# CancerGD: a resource for identifying and interpreting genetic dependencies in cancer

**DOI:** 10.1101/081992

**Authors:** Stephen Bridgett, James Campbell, Christopher J. Lord, Colm J. Ryan

## Abstract

Genes whose function is selectively essential in the presence of cancer associated genetic aberrations represent promising targets for the development of precision therapeutics. Here we present CancerGD (www.cancergd.org), a resource that integrates genotypic profiling with large-scale loss-of-function genetic screens in tumor cell lines to identify such genetic dependencies. CancerGD provides tools for searching, visualizing, and interpreting these genetic dependencies through the integration of functional interaction networks.

## Main text

The ability to inhibit tumors in molecularly defined cohorts of patients is a cornerstone of precision cancer treatment. A successful approach has been the development of drugs that inhibit proteins specifically required in tumors harboring aberrations in recurrently altered cancer ‘driver genes’ [1]. For example, oncogene addiction effects, such as the increased sensitivity of *ERBB2* (*HER2)* amplified breast tumors to *ERBB2* inhibitors [2], can be clinically exploited, as can non-oncogene addiction effects, such as the synthetic lethal relationship between *BRCA1*/*BRCA2* mutations and PARP inhibitors [3]. To identify additional cancer genetic dependencies (CGDs) that may ultimately be exploited therapeutically, multiple groups have performed large-scale loss-of-function genetic screens in panels of tumor cell lines[4–8]. Integrating the results of these screens with molecular profiling data creates hypothesis-generating resources where the hypotheses are of the form *‘tumor cells with a mutation in gene X are sensitive to inhibition to of gene Y’*. These hypotheses are typically tested in subsequent experiments – for example, in larger panels of cell lines, using orthogonal mechanisms of gene inhibition, and/or in mouse models – to ensure they are not statistical or experimental artefacts. Recent examples of novel CGDs discovered through genetic screening approaches include an increased sensitivity of *ARID1A* mutant cell lines to inhibition of the *ARID1A* paralog *ARID1B* [9], of *PTEN* mutant breast tumor cell lines to inhibition of the mitotic checkpoint kinase *TTK* [4], and of *MYC* amplified breast tumor cell lines to inhibition of multiple distinct splicing components [10].

Although the results of loss-of-function screens are typically made publically available, their integration with genotypic data remains challenging for those without bioinformatics skills. Sequencing and copy number data must be processed to identify likely functional alterations, cell line names matched between different data sources, and statistical analysis performed to identify associations between the alteration of driver genes and an increased sensitivity to inhibition of target genes. To address these challenges we have developed CancerGD (www.cancergd.org), a resource that integrates multiple loss-of-function screens [5, 7, 8] with genotype data [11–13] to identify CGDs associated with a panel of cancer driver genes.

CancerGD currently facilitates the searching, visualization, and interpretation of CGDs (Figure 2) associated with 36 driver genes (Supplementary Table 1). These genes were selected based on their identification as driver genes in multiple independent analyses [5, 11, 14] and due to their alteration in at least five tumor cell lines featured in one or more of the included loss-of-function screens. Driver gene associated CGDs are identified both across cell lines from multiple histologies (‘Pan Cancer’) and within tumor cell lines arising from specific primary sites (e.g. ‘Breast’). With an intuitive search interface it is thus possible to retrieve CGDs associated with *ERBB2* amplification across cell lines from all tissue types or specifically associated with *ERBB2* amplification in breast tumor models (Figure 2A). The data supporting every CGD can be visualized in an interactive box plot (Figure 2B) and downloaded for reference.

**Figure 1.**
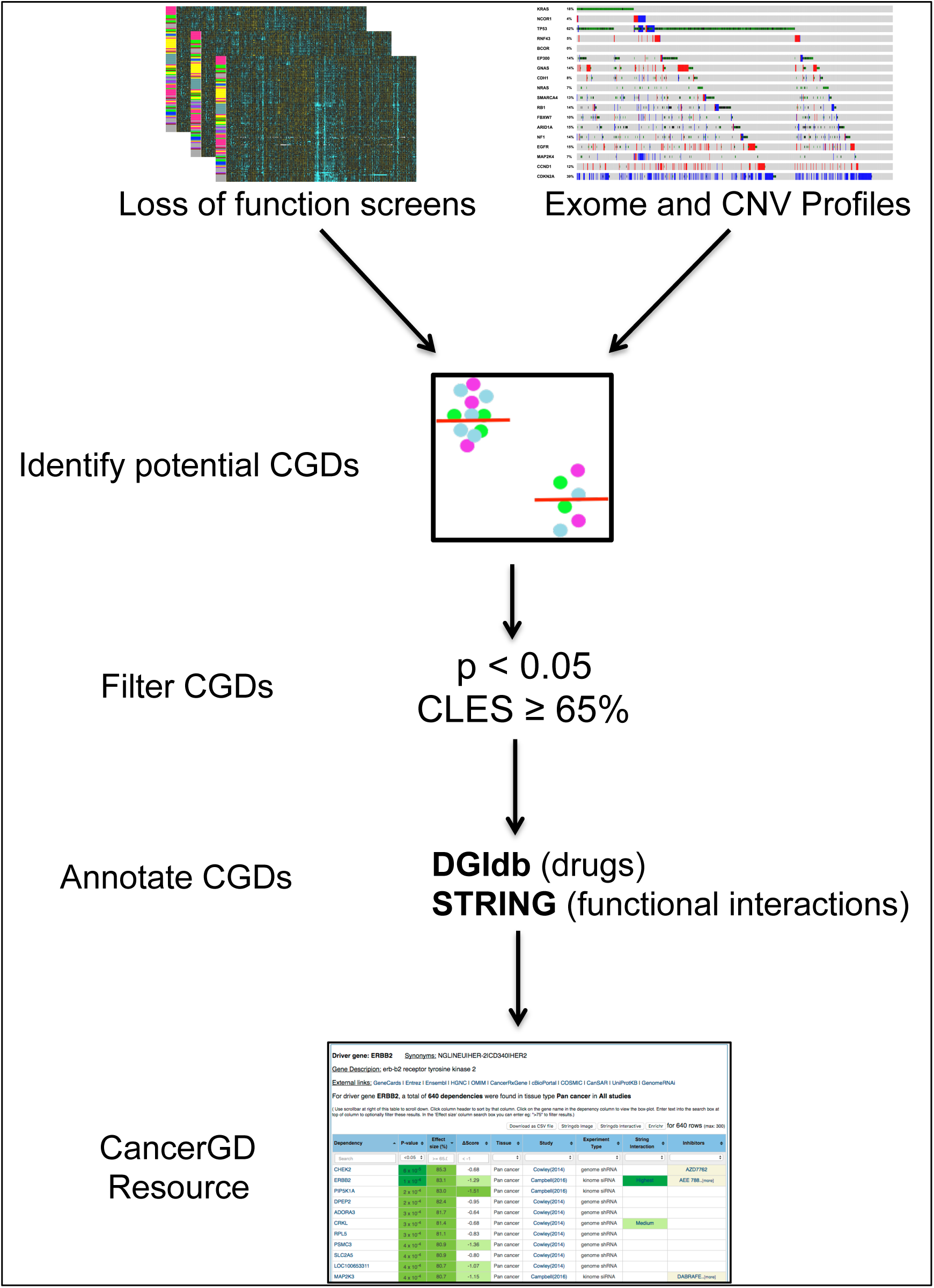
CancerGD overview. Loss-of-function screens from multiple sources are integrated with exome and copy number profiles from the GDSC resource. Cell lines are annotated according to the mutational status of a panel of driver genes. Statistical analysis is then performed to identify associations between the presence of driver gene alterations and sensitivity to reagents targeting specific genes. These CGDs are filtered such that only those with nominal significance (p<0.05) and moderate common language effect sizes (≥ 65%) are retained. Finally all CGDs are annotated according to whether they occur between driver-target pairs with known functional relationships (STRING) and whether there is an inhibitor available for the target gene (DGIdb).

**Figure 2.**
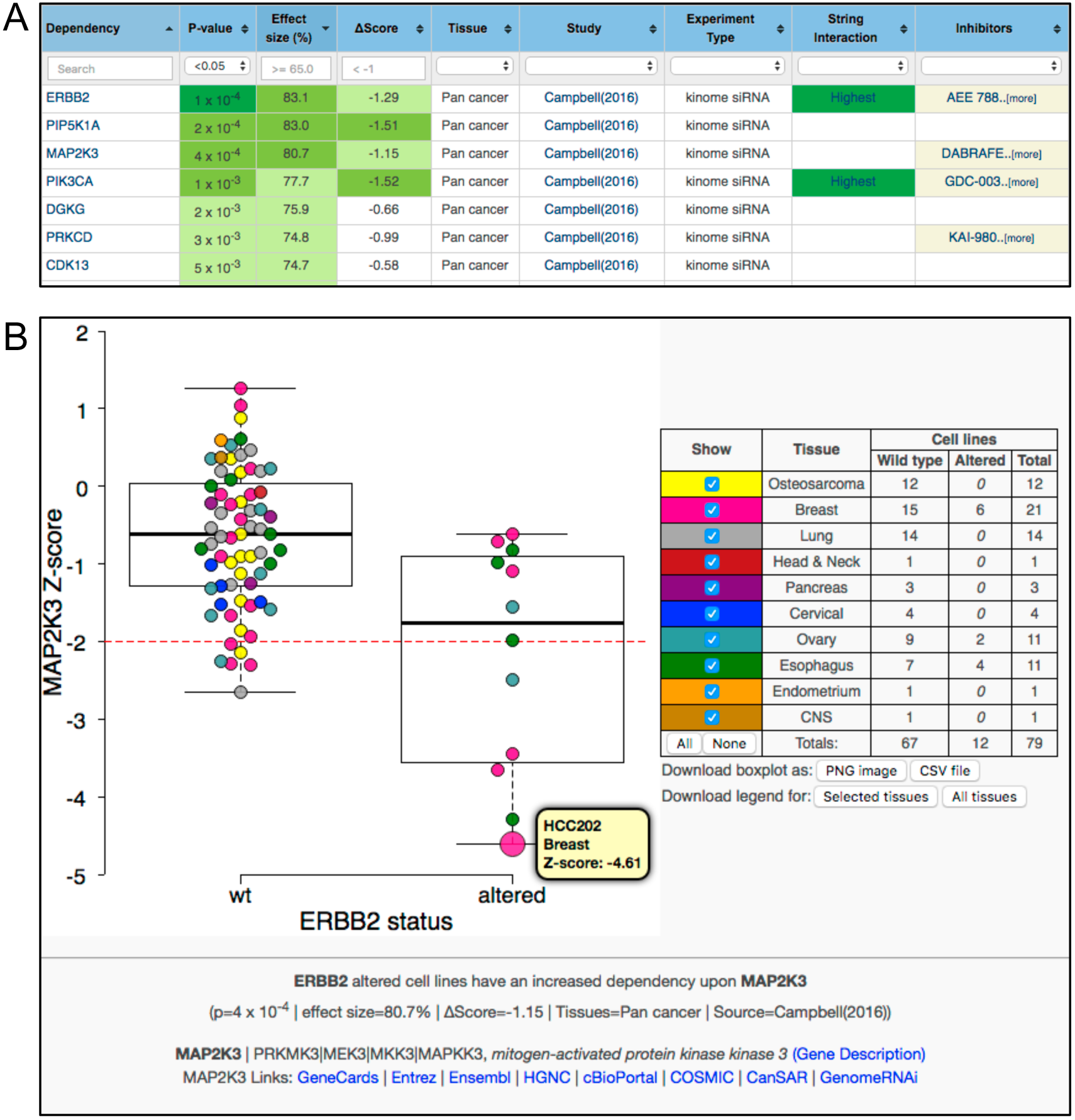
Genetic dependency exploration and visualization. A) The principle view of the database. Each row represents a gene identified as a dependency associated with *ERBB2* amplification in Campbell *et al*[5] across all tumor types (Pan cancer). Columns display experimental details along with the p-value, common language effect size and difference in median sensitivity score for each dependency. Genes with a known functional relationship to the driver gene (e.g. *PIK3CA*) are indicated in the ‘String interaction’ column and drugs known to inhibit the target gene indicated in the ‘Inhibitors’ column. Toggles/search boxes permit easy filtering of interactions – e.g. to select only those genes with an associated inhibitor available. B)Example boxplot showing an increased sensitivity of *ERBB2* amplified cell lines to inhibition of *MAP2K3*. Each circle represents the sensitivity of a particular cell line to RNAi reagents targeting *MAP2K3*. Cell lines are grouped according to *ERBB2* amplification status with the wild-type group on the left and amplified group on the right. Cell lines are coloured according to site of origin and toggles on the right permit the hiding/showing of cell lines from specific sites. Hovering over a given shape provides the cell line’s name, the primary site, and the score associated with the RNAi reagent in that cell line. An overlapped box-whisker plot displays the interquartile range and the median for each group. High-resolution PNG images for each box plot can be downloaded along with a CSV file containing all of the data presented in the box plot. Links to the target gene (*MAP2K3*) on additional sites are provided at the bottom of the plot.

Aside from oncogene addiction effects [1], which represent a tiny minority of the dependencies stored in CancerGD, the mechanistic interpretation of CGDs remains challenging. Why would mutation of one gene result in an increased dependency upon another? In yeast, the interpretation of such relationships has been greatly aided by the integration of protein-protein interaction networks with genetic screens [15]. Following a similar model, to aid the interpretation of CGDs in CancerGD we integrate functional interactions from the STRING database [16]. This facilitates the rapid identification of CGDs involving gene pairs with known functional relationships. For instance in the Campbell *et al* dataset [5] *ERBB2* amplification is associated with an increased dependency upon the *ERBB2* protein interaction partners *JAK2* and *ERBB3*, as well as the *ERBB2* downstream effector *PIK3CA* (Figure 2A). Similarly in the Cowley *et al* dataset [7] loss or mutation of the BAF complex subunit *ARID1A* is associated with an increased dependency upon the *ARID1A* paralog and BAF complex member *ARID1B* [9]. Such dependencies may make more promising candidates for follow on experiments as they are supported by existing functional relationships in addition to the genetic association.

In addition to identifying known functional interactions between the driver gene and associated dependency, it can be helpful to understand the relationships between all of the CGDs associated with a given driver gene. For instance we previously found that cell lines with a deletion or mutation of the tumor suppressor *SMAD4* display a strong dependency upon the mitotic checkpoint kinase *CHEK1* [5]. Considered in isolation it is not clear whether this CGD relates to a specific function of *CHEK1* or a more general sensitivity to inhibition of the mitotic checkpoint. However, by analysing all of the dependencies associated with *SMAD4* we found that they were densely connected on the protein interaction network and primarily involved in the mitotic checkpoint [5], suggesting a more general sensitivity to perturbation of this pathway. To facilitate the identification of such pathway-level dependencies CancerGD provides network visualizations of the functional interactions between CGDs associated with each driver gene (Supplementary Figure 1).

In contrast to the results of drug screening efforts in panels of tumor cell lines [12, 13, 17–20], the CGDs identified in loss-of-function screens include targets that have no inhibitors available and consequently may serve as the rationale for the development of new small-molecule inhibitors. To facilitate the identification of CGDs that may be more readily exploited with available inhibitors CancerGD integrates drug-gene interaction relationships from DGIdb [21].

The loss-of-function screens currently included in CancerGD were all performed using RNA interference (RNAi) approaches. Although existing CRISPR based loss-of-function screens include a relatively small number of cell lines [22–24] it is clear that CRISPR screens in larger panels of tumor cell lines will soon become available. Nothing in the functionality or implementation of CancerGD is specific to RNAi screens and consequently as larger scale CRISPR screens become available we will incorporate their results into the resource.

We believe that CancerGD will be a useful resource to aid a wider group of cancer researchers to benefit from the information generated in large-scale loss-of-function screens.

## Methods

### Genotype data

Exome data for ~1,000 cell lines are obtained from the GDSC resource [12, 13]. We use this data to annotate ~500 driver genes [5] according to whether they feature likely functional alterations. For oncogenes we consider recurrent missense or recurrent in frame deletions/insertions to be likely functional alterations, where recurrence is defined as at least 3 previous mutations of a particular site in the COSMIC database [11]. In addition to recurrent missense or indel events, for tumor suppressors we consider that all nonsense, frameshift and splice-site mutations are likely functional alterations. For copy number analysis we use the gene level copy number scores from COSMIC for the same set of cell lines (which are derived from PICNIC analysis of Affymetrix SNP6.0 array data) [11–13, 19]. An oncogene is considered amplified if the entire coding sequence has 8 or more copies while a tumor suppressor is considered deleted if any part of the coding sequence has a copy number of 0 as per Garnett *et al* [19]. For the majority of driver genes we integrate the two sources together. For all tumor suppressors we consider a functional alteration to be either a deletion (derived from copy number profiles) or a presumed loss-of-function mutation (as identified in the exome data). For most oncogenes we consider a functional alteration to be either an amplification or a recurrent mutation/indel. For a small number of oncogenes (*ERBB2, MYC, MYCN*) we consider only amplifications as functional events, while for another group (*KRAS, BRAF, NRAS, HRAS*) we only consider recurrent mutations/indels.

### Loss of function screens

Three large-scale RNAi datasets are currently included in CancerGD [5, 7, 8]. These include a kinome focussed siRNA screen covering a panel of 117 cell lines from diverse histologies [5], a genome-scale shRNA screen focussed on 77 breast tumor cell lines [8] and a large-scale shRNA screen covering 216 cell lines from diverse histologies [7]. Cowley *et al* [7] is largely a superset of a previous screen from the same lab [6] and hence the two resources are not included separately. Similarly the kinome siRNA screen from Cambell *et al* [5] contains the majority of the breast tumor cell lines screened in a previous breast cancer kinome siRNA screen from the same lab [4] and hence they are not included separately.

### Cell line naming

Internally we follow the naming convention established by the Cancer Cell Line Encyclopedia [17]. The CCLE naming convention is the cell line name (containing only numbers and upper case letters) followed by an underscore, followed by the tissue/primary site in upper case. The cell line names are taken from [12], converted to uppercase and punctuation removed. Where possible we use the same tissue types as the CCLE, in a small number of cases where a tissue was absent from the CCLE (e.g. CERVIX) we have created a new tissue type. Having the tissue type in the cell line name facilitates filtering the boxplots (e.g. to show the gene inhibition sensitivities for cell lines from a specific tissue) in the browser without having to perform additional database queries. Furthermore two of the published loss-of-function screens already follow this naming convention [5, 7] while the third features only breast cell lines and was trivially converted [8]. In instances where the same cell line is featured in two datasets but there is a naming disagreement (e.g. H1299_LUNG in Campbell *et al* [5] is NCIH1299_LUNG in our genotype set) we manually rename the RNAi dataset to match the genotype data.

### Gene identification

CancerGD provides links to multiple external sources that use a variety of different gene identifiers. Consequently for each gene in the database we store multiple identifiers (Entrez Gene ID, Ensembl Gene identifiers, HUGO Gene Names, Ensembl Protein IDs). We also store synonyms for each gene to facilitate easy gene look up (e.g. *ERBB2* can be identified by searching for *HER2*). These synonyms are obtained from the HGNC resource [25].

### Drug target annotations

Drug-gene relationships are obtained from the Drug-Gene Interaction Database (DGIdb), which integrates drug-gene relationships from multiple sources [26]. Only inhibitor relationships are retrieved, as we are interested in drugs that inhibit the products of specific genes, rather than drugs whose efficacy is associated with the mutation of specific genes. Results from DGIdb sourced from MyCancerGenome and MyCancerGenomeClinicalTrial are excluded for the same reason.

### Statistical analysis

We use R for all statistical analysis. For each driver gene / target gene combination we compare cell lines harbouring a likely functional alteration in the driver gene to cell lines with no alteration in that gene and test if the cell lines with the functional alteration are more sensitive to RNAi reagents that inhibit that gene. This is tested using a one-sided Mann-Whitney U test. A variety of alternative two-sample tests have been used in previous publications, including median permutation tests [4, 5] and mutual information based measures [7]. The Mann-Whitney U test has a number of advantages for CancerGD – it is rapid to calculate and it does not assume that the scores for each gene are normally distributed. The latter is important as it means the test can be used uniformly on loss-of-function screens from multiple sources that use different scoring schemes. For all screens we use the authors’ provided scoring scheme (zGARP for Marcotte *et al* [8], ATARIS phenotype score for Cowley *et al* [7], and robust Z-score for Campbell *et al* [5]). In addition to the p-value from the Mann-Whitney U test we calculate a common language effect size (CLES) for each dependency. The CLES is equivalent to the *Area under the ROC curve* and the *Probability of Superiority* and indicates the probability that a cell line with an alteration in a particular driver gene is more sensitive to a given RNAi reagent than a cell line without that alteration. In the database we store all nominally significant dependencies (p<0.05) with a CLES ≥ 0.65. In a small number of instances multiple ATARIS scores are presented for a single gene – when storing CGDs we incorporate the ATARIS score with the lower p-value.

### Functional interactions

Functional interactions are obtained from STRING. We store all interactions that are medium confidence (STRING score > 0.4) or higher. Cut-offs to identify interactions as ‘Medium’, ‘High’ and ‘Highest’ confidence are those defined by STRING. For displaying the functional interactions between the dependencies associated with each driver gene we use the STRING API [16].

### Implementation

CancerGD is implemented in Python using the Django framework and follows a model/view/controller architecture. JQuery is used for Javascript processing in the browser interface. MySQL is used by default for data storage but SQLite can be used for development / testing purposes with minimal documented changes. The application is currently hosted on the PythonAnywhere system, a generic Python web services host, suggesting that the application is portable.

### Code availability

Source code for the entire project (R/Python/Javascript/HTML) is publicly available on GitHub (https://github.com/cancergenetics/cancergd). Detailed instructions on how to run the statistical analysis, install the web application and populate the database are also provided in the GitHub repository (CancerGD_Manual_v1.1.doc).

## Author Contributions

SJB wrote code for the database. JC contributed R code for statistical analysis. CJL provided guidance on the design of the database and the manuscript. CJR conceived and designed the resource and wrote the manuscript. All authors read and approved the final manuscript.

## Acknowledgements

We thank Lars Juhl Jensen for help with STRING integration. We also thank members of the ICR Gene Function Team and Systems Biology Ireland for providing feedback on earlier versions of this resource. Work on this project was supported by a grant to CJR from the Irish Health Research Board (KEDS-2015-1636). CJR is a Sir Henry Wellcome Fellow jointly funded by Science Foundation Ireland, the Health Research Board, and the Wellcome Trust (grant number 103049/Z/13/Z) under the SFI-HRB-Wellcome Trust Biomedical Research Partnership. CJL is supported by Cancer Research UK (grant number C347/A8363) and Breast Cancer Now. We acknowledge NHS funding to the ICR/Royal Marsden Hospital Biomedical Research Centre.

---

**Supplementary Figure 1.**
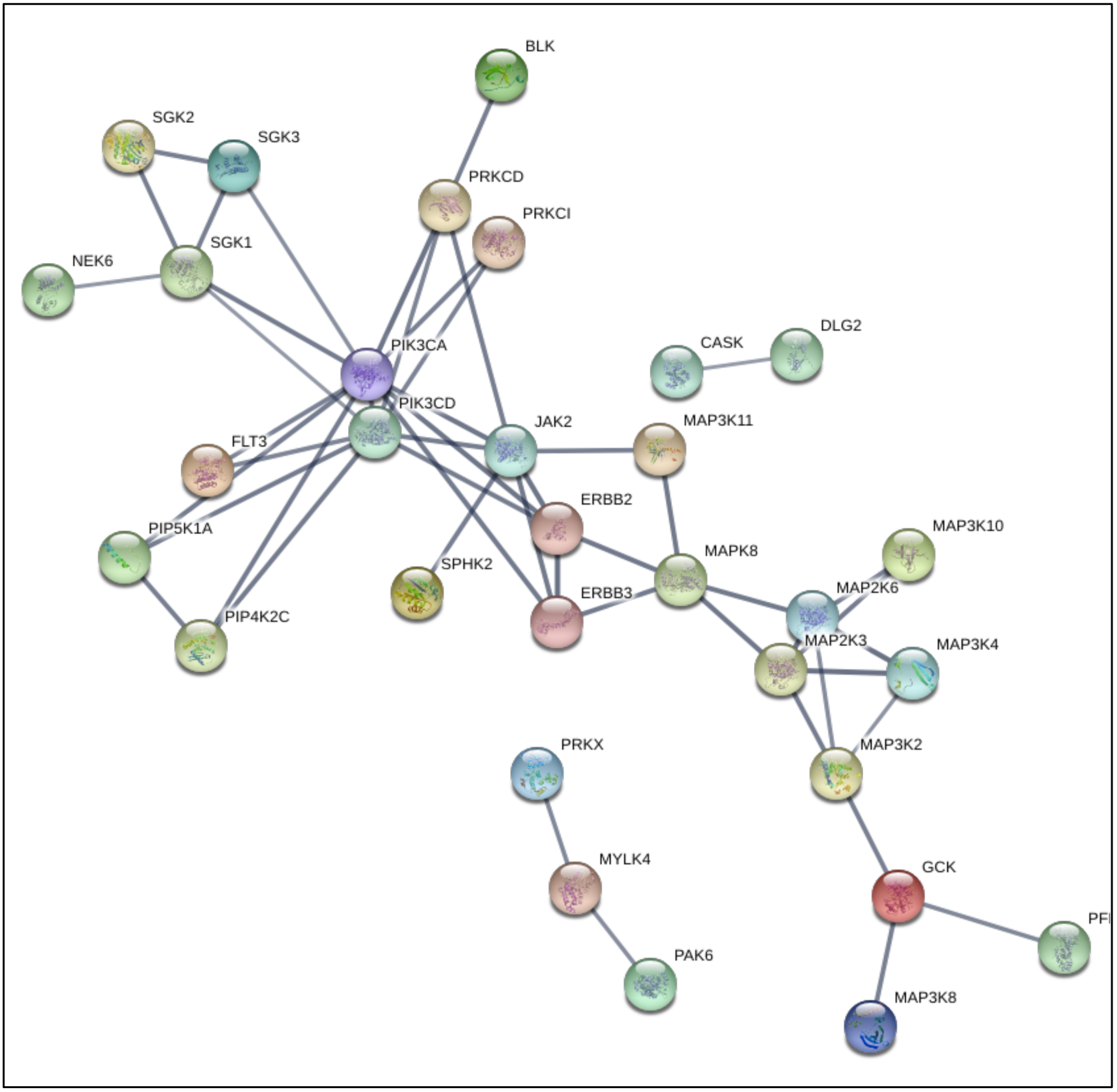
Visualizing the interactions between all CGDs associated with a specific driver gene. High confidence STRING functional interactions between CGDs associated with *ERBB2* amplification in Campbell *et al* are shown.

**Supplementary Table 1. Driver genes currently included in CancerGD**

